# Multistability in neural systems with random cross-connections

**DOI:** 10.1101/2023.06.05.543727

**Authors:** Jordan Breffle, Subhadra Mokashe, Siwei Qiu, Paul Miller

**Affiliations:** Neuroscience Program, Brandeis University, 415 South St, Waltham, MA 02454; Volen National Center for Complex Systems, Brandeis University, 415 South St, Waltham, MA 02454; Department of Biology, Brandeis University, 415 South St, Waltham, MA 02454; Department of Neurology, Cedars-Sinai Medical Center, Los Angeles, CA, USA

**Keywords:** Attractor basin, mean field, fixed points, bistable

## Abstract

Neural circuits with multiple discrete attractor states could support a variety of cognitive tasks according to both empirical data and model simulations. We assess the conditions for such multistability in neural systems, using a firing-rate model framework, in which clusters of neurons with net self-excitation are represented as units, which interact with each other through random connections. We focus on conditions in which individual units lack sufficient self-excitation to become bistable on their own.

Rather, multistability can arise via recurrent input from other units as a network effect for subsets of units, whose net input to each other when active is sufficiently positive to maintain such activity. In terms of the strength of within-unit self-excitation and standard-deviation of random cross-connections, the region of multistability depends on the firing-rate curve of units. Indeed, bistability can arise with zero self-excitation, purely through zero-mean random cross-connections, if the firing-rate curve rises supralinearly at low inputs from a value near zero at zero input. We simulate and analyze finite systems, showing that the probability of multistability can peak at intermediate system size, and connect with other literature analyzing similar systems in the infinite-size limit. We find regions of multistability with a bimodal distribution for the number of active units in a stable state. Finally, we find evidence for a log-normal distribution of sizes of attractor basins, which can appear as Zipf’s Law when sampled as the proportion of trials within which random initial conditions lead to a particular stable state of the system.

## 1. Introduction

An extensive literature in neuroscience suggests that neural activity can proceed through sequences of distinct states during sensory processing, motor output, or memory-based decision making (Abeles et al., 1995; Benozzo et al., 2021; Escola et al., 2011; Jones et al., 2007; La Camera et al., 2019; Mazzucato et al., 2015; Miller, 2016; Morcos & Harvey, 2016; Rainer & Miller, 2000; Seidemann et al., 1996). The distinct states are revealed as patterns of neural activity that remain relatively stable for durations much longer than those of the rapid transitions between states. Models of the underlying circuitry assume the states correspond to fixed points (or the remnants of fixed points) of the system (Ballintyn et al., 2019; La Camera et al., 2019; Mazzucato et al., 2019; Miller, 2013; Miller & Katz, 2010; Rabinovich et al., 2001; Rabinovich et al., 2014; Recanatesis et al., 2022; Taylor et al., 2022) with the itinerancy from fixed point to fixed point known as latching dynamics (Boboeva et al., 2021; Lerner et al., 2012, 2014; Lerner & Shriki, 2014; Linkerhand & Gros, 2013; Russo & Treves, 2012; Song et al., 2014; Treves, 2005).

Transitions between fixed points can be due to their inherent instability when they are saddle points. Otherwise, in networks where a reduced model of the system possesses multiple stable fixed points, transitions arise from one or more of (1) an external stimulus, (2) noise fluctuations, or (3) the drift of a slow variable which impacts a parameter in the reduced model causing it to cross a bifurcation point. Since the number of stable fixed points becomes a key indicator of the potential information processing or memory capacity of the network, it is important to understand the conditions under which a system possess multiple stable fixed points.

Here we use firing-rate models (Wilson & Cowan, 1973), in which each unit represents a cluster or assembly of similarly responsive neurons with stronger connections within each cluster as observed in some cortical circuits (Perin et al., 2011; Song et al., 2005). Such assemblies can arise in response to a lifetime of stimuli via Hebbian plasticity (Hebb, 1949), which increases connection strengths between excitatory neurons with correlated activity (Bourjaily & Miller, 2011; Brunel, 2003). We assume random interactions between such clusters (Stern et al., 2014), representing the result of a history of uncorrelated stimuli.

Each isolated stable fixed point in a system is an attractor state, with a basin of attraction determined by the set of initial conditions that result in neural activity settling at (after being “attracted to”) the fixed point. Systems with many such attractor states have provided the framework for understanding pattern completion and separation of new inputs following memory encoding of stimuli, since the highly influential work of Hopfield and others (Anishchenko & Treves, 2006; Battaglia & Treves, 1998; Hopfield, 1982; Hopfield, 1984; Treves, 1990; Zurada et al., 1996). Indeed, there is abundant evidence of such attractor states in neural circuits (Daelli & Treves, 2010; Fuster, 1973; Goldberg et al., 2004; Golos et al., 2015; Wills et al., 2005), perhaps most obvious to us when an ambiguous stimulus can cause perceptual alternation due to activity flipping between two (quasi-stable) attractor states (Moreno-Bote et al., 2007). However, while the number of stable states in systems such as the Hopfield network (Hopfield, 1982; Hopfield, 1984) have been characterized (Amit et al., 1985a, 1985b; Folli et al., 2016), the connections between units in such networks are correlated (in fact, the connectivity matrix is symmetric), so it is unclear to what extent multiple attractor states would arise in a nonsymmetric random network.

Work by others (Stern et al., 2014) showed that when each unit has sufficient self-excitation to become bistable (and therefore become in essence a memory element in of itself) multiple attractor states are possible in a network with non-symmetrically randomly connected units. Such a result is trivial in the limit of zero cross-connection strength, in which case a system of *N* bistable units possesses 2^*N*^ stable states. In the randomly connected system, studied in the large-*N* limit, increased strength of random cross-connections decreases the number of multistable states, eventually rendering the system chaotic as all fixed points become unstable. With weaker self-connections, the network would be either quiescent or, given sufficient cross-connection strength, chaotic (Sompolinsky et al., 1988).

Here we find that such results depend on the form of the input-output function (the firing-rate, or f-I curve) of a neuron. Indeed, if we assume neurons have low firing rates in the absence of input, random non-symmetric cross-connections can lead to multistability, even when individual units have zero self-excitation.

In the following sections, we first present simulations showing the types of activity possible and their observed coexistence in networks of up to 1000 randomly coupled units. We then show the phase diagrams in the large-*N* limit of such systems. Finally, we present results for systems with binary activation functions, for which we develop an alternative mean-field analytic approach that we use for finite-as well as infinite-*N* systems. Also, given the more rapid simulations when activations are binary, we provide a more thorough analysis of the attractor states in such systems.

## 2. Simulations of Finite Networks

We simulated networks of *N* randomly connected firing rate units with response function *f*(*x*) representing their output to total input, *x*. The total input, *x*_*i*_, to the *i*-th unit is described by the dynamical equation with time constant, *τ*:

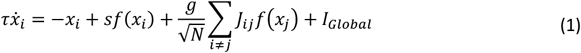

where *s* and *g* are parameters that scale the self-connection and cross-connection strengths, respectively, and *J*_*ij*_ is the matrix of normalized cross-connection strengths drawn from a normal distribution with zero mean and unit variance. *I*_*Global*_ is a constant input that inhibits or excites the whole network (equivalent to a shift in threshold, *x*_*th*_) and is kept at zero unless stated otherwise.

We simulated models with distinct single-unit response functions, *f*(*x*), in order to assess its role in network dynamics:

1. Hyperbolic tangent: *f*(*x*) = tanh(*x*) and our model is identical to that of (Stern et al., 2014). We also compare more general forms, 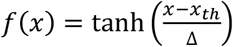 to connect results to those of the logistic function with less symmetry in the firing rates (*i*.*e*., units require net excitatory input to reach half their maximum rate if *x*_*th*_ > 0).
2. Logistic function: 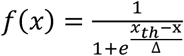 where *x*_*th*_ is a threshold input required for the firing rate of a unit to reach half of its maximum value and Δ is inversely proportional to the steepness of the response function.
3. Binary output via the Heaviside function: *f*(*x*) = *Heaviside*(*x* − *x*_*th*_), which is equivalent to the logistic function in the limit Δ→ 0.

In all systems we adjusted the threshold parameter, *x*_*th*_, for a given steepness of response function (*i*.*e*., a given value of Δ), such that in the absence of cross-connections (*g* = 0) the system becomes multistable, because each unit is bistable, at *s* = 1. This allows us to compare results across systems with different single-unit response functions, *f*(*x*) (Figure 1, see Appendix A).

**Figure 1.**
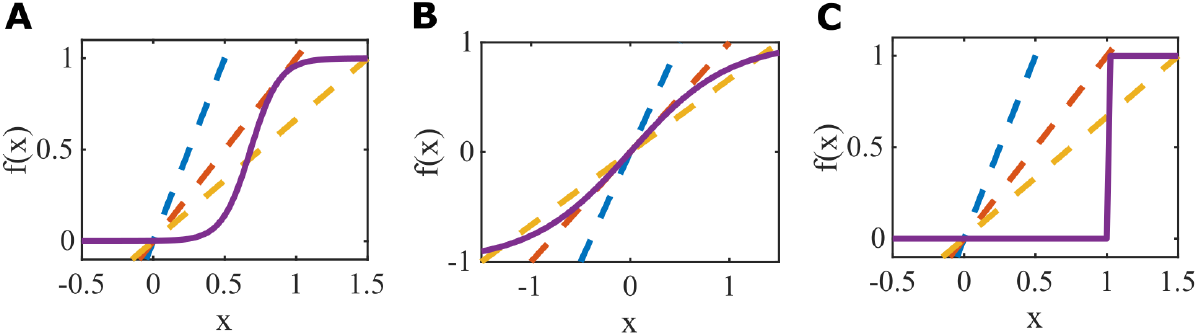
Single-unit response functions, *f*(*x*), produce bistability at *s* = 1. A. Logistic function shown with slope parameter, Δ= 0.1, and threshold, *x*_*th*_ = 0.681. **B**. Tanh function with Δ= 1, and threshold, *x*_*th*_ = 0. **C**. Heaviside function with binary response, equivalent to the logistic function with Δ= 0, and *x*_*th*_ = 1. Feedback curves, *I* = *sf* shown as dashed lines with blue *s* = 0.5, red *s* = 1, and yellow *s* = 1.5 demonstrate the bifurcation from an inactive state to bistability at *s* = 1. Note that the bifurcation is a saddle-node in **A** and **C** but a pitch-fork in **B**.

### 2.1. Observed forms of simulated network dynamics with logistic function responses

We define the network state by its long-term activity, which can be either constant, oscillating, or chaotic. We separate out constant (stable) states into two types: those that include active units and those with only inactive units (and are quiescent). We therefore obtain four labels for final states: quiescence, stable activity, limit cycle, and chaos.

In many networks we find, by varying initial conditions, the existence of more than one type of state in a single network. Some networks have multiple forms of all four activity types, such as the example network in Figure 2A-D. This example network of logistic units (*N* = 100, *s* = 0, *g* = 3.25, and Δ = 0.1) has a stable quiescent state, two stable active states, two unique stable limit cycles, and a chaotic attractor. Example trials leading to each of these distinct states in the same network are shown as a subset of units’ firing rates (Figure 2A) and in principle component space (Figure 2B). These states have similar root mean squared (RMS) firing rates, except for the quiescent state (Figure 2C).

**Figure 2:**
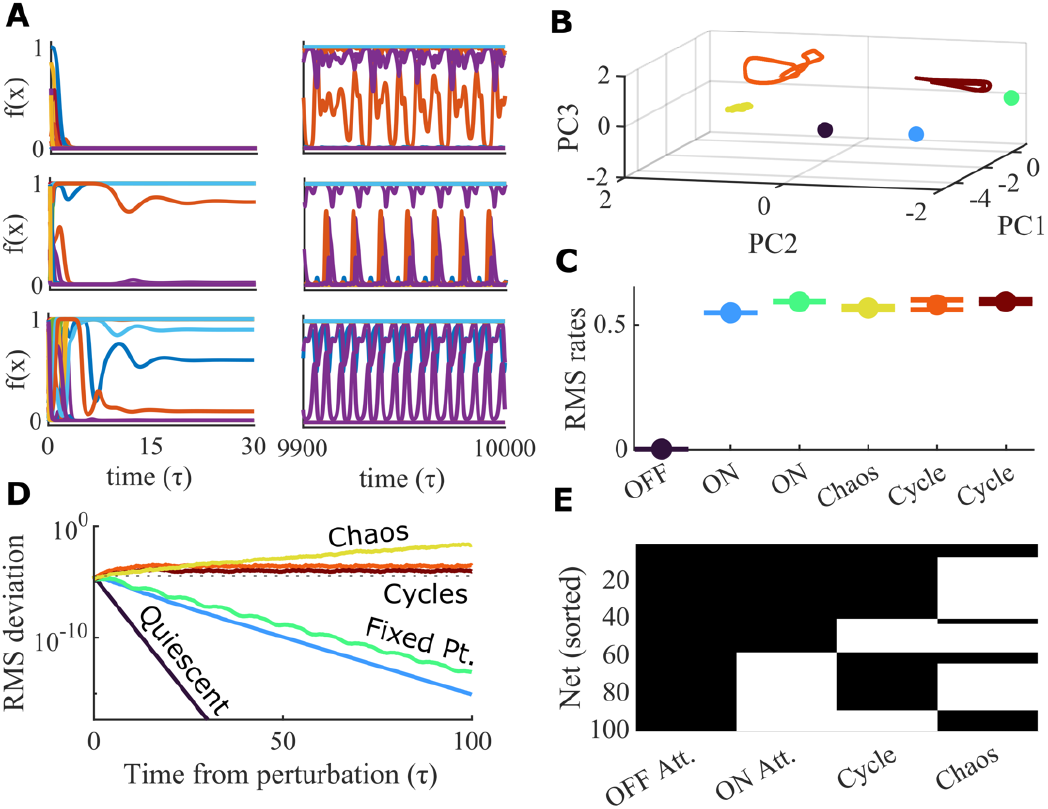
All activity regimes can be observed in a single random network. **A**. Different initial conditions in the same network can lead to a quiescent state, two stable active states, two stable limit cycles, and chaotic activity. For each example condition, the firing rates of a subset of units is plotted for either the beginning or end of the trial. The parameters for this network are *N* = 100, *s* = 0.0, *g* = 3.25, and *Δ* = 0.1, with a logistic FI curve. **B**. The first three principal components from PCA of the firing rates from the trials in **A**. Each color is a different trial. All trials converge to one of the six different activity regimes show in **A**. The last 100 *τ* of each trial is plotted to show the steady state activity. **C**. The root mean squared (RMS) firing rates of the network for each of the 6 example simulations. Colors correspond to those in **B**. The mean and standard deviation of the last 100 *τ* of each simulation is plotted. “OFF” = quiescent state. “ON” = stable active state. **D**. For each trial 100 perturbations of it were simulated and the RMS deviation from the unperturbed simulation of the firing rates of all units in the network was calculated across the time from perturbation. Colors correspond to the trial colors in **B** and **C**. Dashed line indicates the initial perturbation magnitude. The median value of the 100 perturbations is plotted. **E**. A diverse array of mixed activity regimes is found across 100 random networks for the same parameters as the example network in **A**-**D**. Black indicates that the network had at least one out of 100 trials with the indicated activity type. All 100 networks had at least two types of activity. Seven out of the 100 networks had all four activity types (the top seven rows).

Perturbation analysis confirms the classification of each of the trials in Figure 2A-C (Figure 2D). For each trial, we simulated 100 perturbations and calculated the median RMS deviation of the perturbed simulation from the original simulation. Analysis of the cross connections of the example network in Figure 2A-D shows that it is not an outlier from an expected random network (Supplemental Figure 1), suggesting this combination of mixed activity states may be a common occurrence. We performed the same perturbation-based classification of activity states for 100 different random networks at the same parameter values and found that all networks show at least two forms of activity (Figure 2E).

Of particular interest are systems without self-connections, (*s* = 0, such as that shown in Figure 2), for which there is a well-established single transition from quiescence to chaos at *g* = 1, when *f*(*x*) = tanh (*x*) (Sompolinsky et al., 1988). When, instead, we use the logistic function, 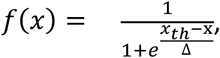, which is simply a scaled and shifted transformation of the tanh function to non-negative values of firing rate, we find a richer set of states in our simulations. Perhaps surprising, circuits without self-connections can exhibit multiple stable states: sometimes having only a low activity state (a quiescent state) with a state of higher net activity (an active state) such as the example network in Figure 3A.

**Figure 3:**
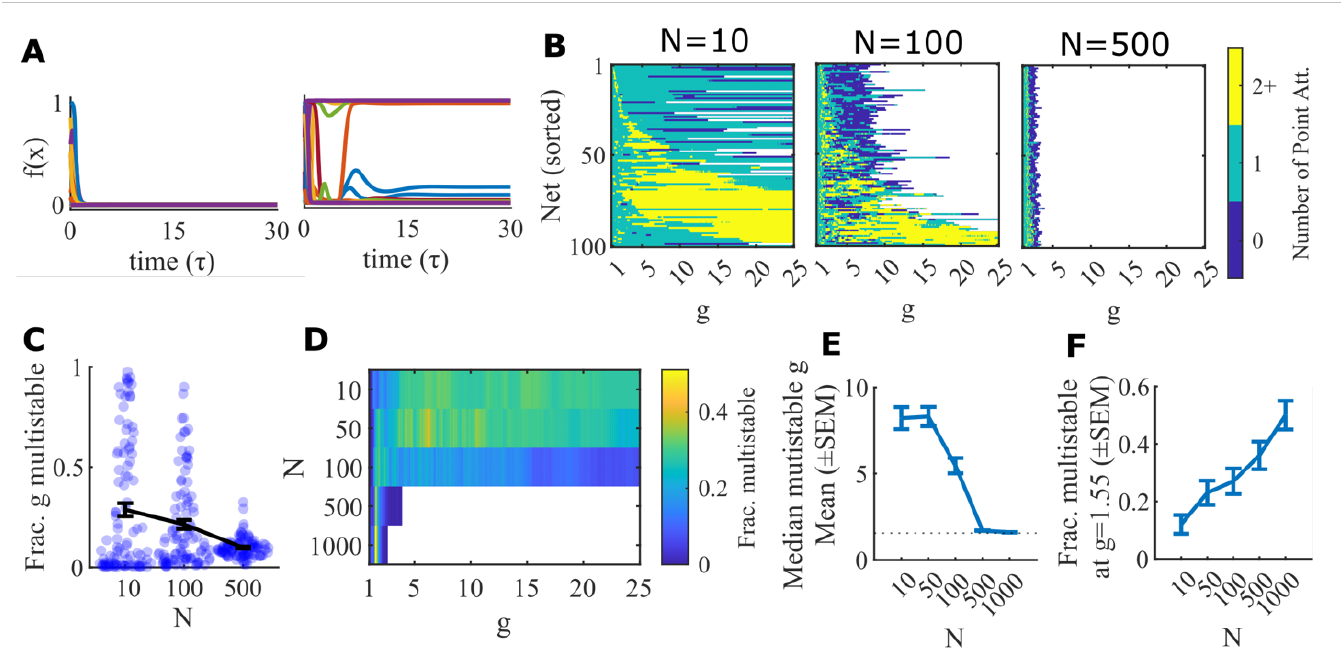
Smaller networks are multistable at larger g values. **A**. An example network (logistic units, *Δ* = 0.2, *N* = 100) that has only the quiescent attractor and one active point attractor, even with no self-connections (*s* = 0, *g* = 2.0). Left, a subset of units’ firing rates in a trial that converges to the quiescent attractor. Right, a trial that converges to an active point attractor. **B**. 100 random networks of logistic units (*Δ* = 0.2) with no self-connections (*s* = 0) of varying size (*N* = 10, 100, 500) were simulated across *g* values. The same network can gain and lose mulistability as g is scaled. Color scale indicates number of point attractors found within 100 trials. White indicates *g* values that were not simulated due to computational limits, wherein each network was allotted 24 hours of cpu time. Networks tended to reach their limit after a series of g values in which the network failed to converge to a fixed point on any trial (shown in blue), indicating the likely end of the networks’ region of multistability. **C**. The fraction of the simulated g values at which each network in **B** was multistable. Bars show mean and SEM. **D**. Fraction of networks that were multistable across the tested *g* values for *N* = 10, 50, 100, 500, 1000. **E**. The median g value at which each network was stable. This converges to *g* = 1.55 (dotted line) as network size increases. **F**. Fraction of networks that are multistable at *g* = 1.55 increases with N.

Indeed, we find that for a given random instantiation of the connectivity matrix, as we scale all connections by *g*, there is, for all 500 total networks tested in Figure 3, some range of connection strengths for which the network is multistable. Supplemental Figure 2 shows how dynamics beyond the existence of multistability changes as *g* is scaled.

We wondered whether such states were the results of a finite size effect, so varied the size of the network (changing *N*). We found that the range of *g* over which we see such multistability at *s* = 0 narrows with increased *N* (Figure 3C), and converges to the same set of values centered on *g* = 1.55 (Figure 3D-F). These and other results prompted us to investigate the phase space of the corresponding infinite-*N* systems via mean field theory and stability analysis (Section 3).

### 2.2. Phase diagram from simulations of finite networks

To assess the likelihood of systems reaching a given type of state across phase space, we simulated 100 networks for each given set of parameters and commenced simulations of each network from 200 initial conditions. Full details of simulation methods are provided in Appendix 1.

In Figure 4 we show that systems with 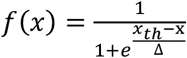 can have multistability at *s* < 1, such that increasing the cross-coupling strength, *g*, increases the fraction of multistable networks for large ranges of *s* and *g*. The observed multistability at low *s* that arises with increasing *g* is most apparent in binary systems (logistic *f*(*x*) with Δ = 0, *x*_*th*_ = 1) and is not so apparent for systems with *f*(*x*) = tanh (*x*). As might be expected, multistability for the logistic function with a steeper slope (Δ = 0.1) is more similar to that of the binary function, and the logistic function with the shallower slope (Δ = 0.2) is more similar to the tanh function.

**Figure 4:**
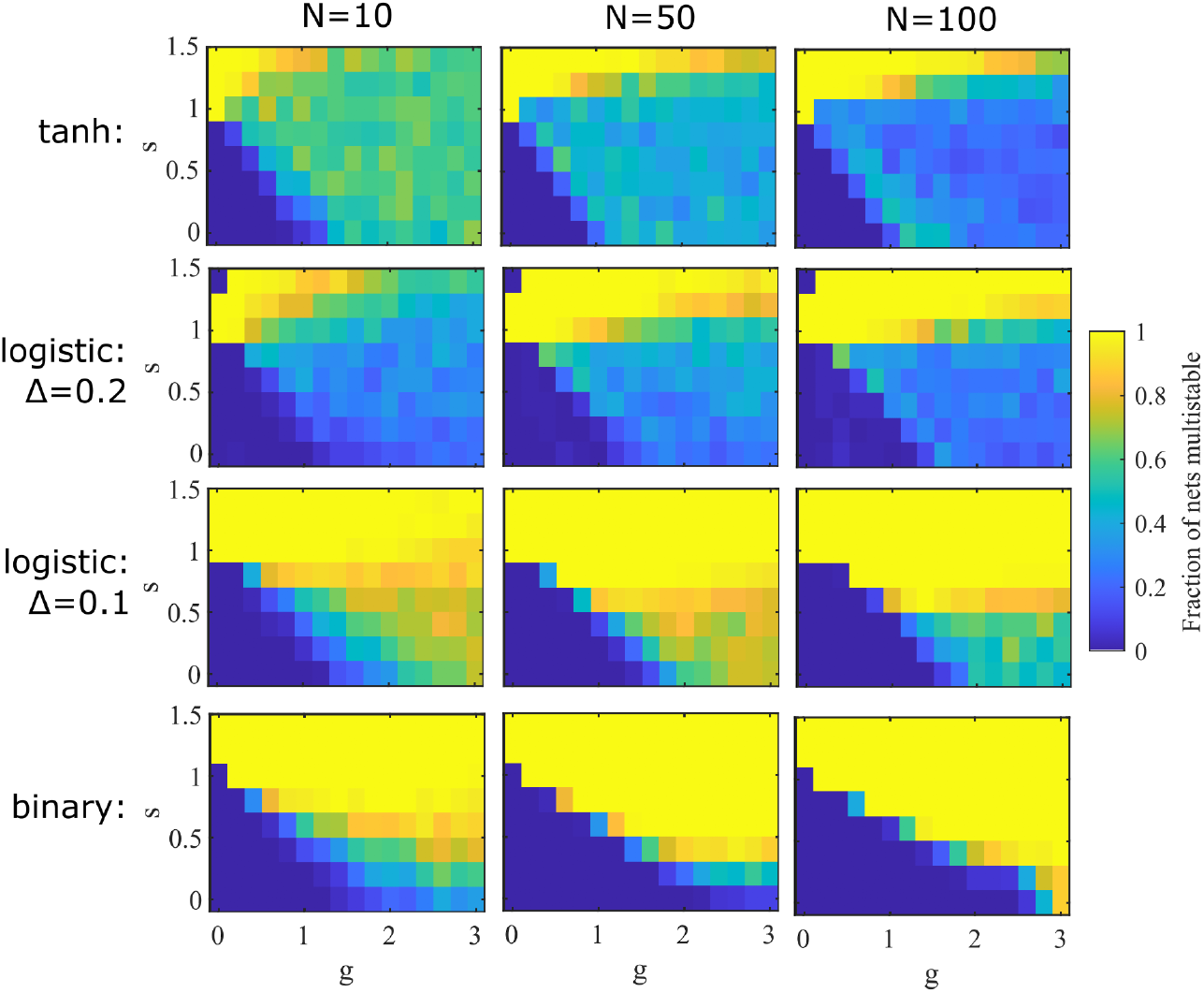
Multistability across phase space. Simulation results showing the fraction of networks at different values of *s* and *g* with multistability. For each parameter point 100 random networks were simulated for 200 random initial conditions. Networks with 10 (left), 50, (center), and 100 (right) units were simulated with the tanh (top), logistic (middle), and binary (bottom) FI curves.

While the impact of the response function, *f*(*x*), on the results in Figure 3 suggest our findings arise from more than a finite size effect, we wanted to test that possibility further. Therefore, we assessed, for different networks with *s* < 1, how the probability of multistability depends on network size. Our goal is to see how reliably cross-connections whose random strengths have a mean of zero could, with increasing standard deviation, *g*, generate multistability that is absent with low *g*.

Simulation results of Figure 3 suggest a peak as a function of network size, *N*, in the likelihood of reaching multiple final states from a fixed (large) number of initial conditions for some parameters. However, without the exhaustive sampling of initial conditions, which becomes unfeasible at large *N*, our lack of multistability at large-*N* is not conclusive of its absence. To proceed further, we calculate results for the infinite-*N* system in Section 3 and develop analysis of binary networks that allows for finite-*N* approximations in Section 4.

### 2.3. Distribution of size of basins of attraction

A major goal in our simulations of neural circuits was to assess the number of stable states they contained as a marker of their information-carrying capacity. In systems with zero cross-connection and strong enough self-interaction (*s* > 1) that each unit could be independently bistable, the number of states is trivially 2^*N*^, the maximal possible for our systems. However, without cross-connections, the ability of such circuits to process information in a history dependent manner vanishes. Also, given the unlikeliness that neural circuits operate in a regime with distinct bistable units, our focus was on circuits with *s* < 1 such that individual units were not bistable, but with sufficiently strong *g* > 0 such that the network could be multistable. For the results in this subsection, we focus on such systems with binary units, *f*(*x*) = *Heaviside*(*x* − *x*_*th*_), with *x*_*th*_ = 1.

For small circuits, *N* < ∼25, one can sample initial conditions systematically with each of the 2^*N*^ combinations of high/low activity per unit tested, but for larger networks one can sample only a subset of all the possible initial conditions. We find that some states have vastly more initial conditions reaching them compared to others. Indeed, if we take the number of randomly chosen initial conditions that reach a particular fixed point as an indication of the size of the corresponding basin of attraction, we find what appears to be a power law (Figure 5). Such a result suggests that in the absence of exhaustive sampling, we will inevitably miss some of the smallest basins of attraction (note that the frequency of visits ranges over 5 orders of magnitude in Figure 5A).

**Figure 5.**
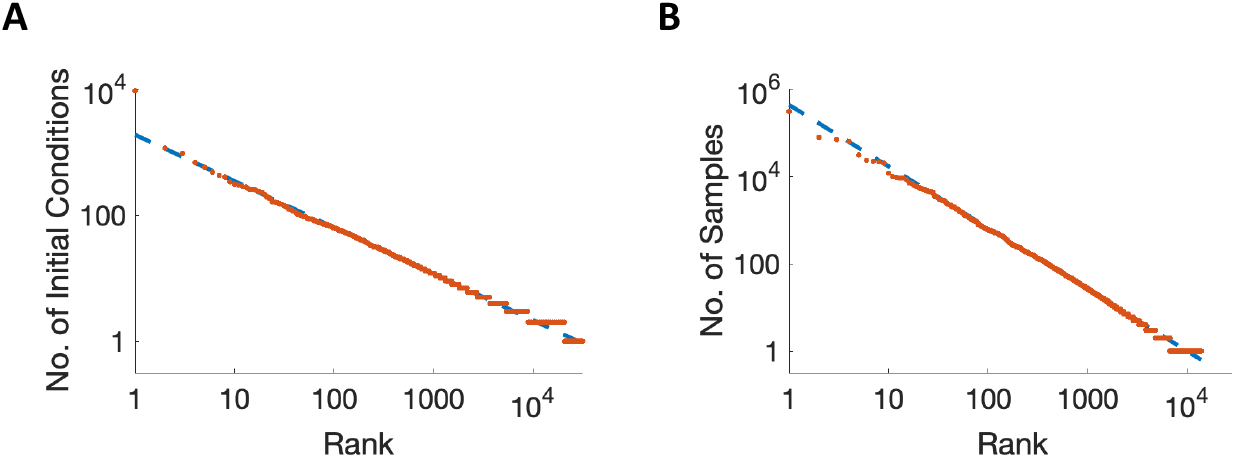
Zipf-like behavior is suggested by the data but likely explained by a log-normal distribution. **A**. The numbers of initial conditions leading to each discovered attractor state (indicating size of basin of attraction) are recorded in an example network of size *N* = 200, with results ordered on a log-log scale. **B**. Simulations of random sampling from a log-normal distribution of sizes of attractor basins, as suggested by analysis, can account for the observed linearity on the log-log scale.

The suggestion of Zipf’s Law in Figure 5A led us to consider theoretical reasons for producing such a distribution. Our conclusion is that the apparent power-law is an artefact produced by sampling a log-normal distribution with a very large width. Our reasoning is as follows. For any stable state some units can have a level of input such that the unit could be stably active or inactive (assuming no change in input from others). If *N* is large, the switching of such a unit does not strongly change the net input to all other units, so another stable state is reached. Such a switch to a different state indicates the crossing to a different basin of attraction. Across all of the distinct states in the network, the number of units with inputs in the bistable range allowing for such stable switching is distributed as a Binomial (if we ignore correlations), which is approximately a Normal distribution at large *N*. Or, equivalently, the number of units that can be switched in any state *without* changing to a different basin of attraction follows a Normal distribution across states. Additionally, the number of combinations of switching a unit without producing a new stable state is approximately exponential in the number that can be individually switched. Combining the two heuristics would suggest a log-normal distribution of sizes of basins of attraction.

To test if our results in Figure 5A were compatible with a log-normal distribution, we generated 10^2^ samples from a fictitious system with 10^2^ states whose sizes, *x*, were distributed as 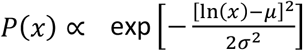, with *μ* = 30 and σ = 6. Such a system was chosen to resemble the statistics, in terms of numbers of states selected and maximum number of selections for any network of one of our random systems with *N* = 200.

The results of our random sampling of a fictitious log-normal system in Figure 5B, indicate that the observation of Zipf’s Law from sampling basins of attraction is, indeed, compatible with a log-normal distribution of attractor-basin sizes. The reason being that many of the small basins, whose expected number of visits is less than one, either do not appear in the sampling at all, or appear once or twice, and therefore increase the number of low-frequency states in a manner that “straightens out” the inverted parabola that would be seen following an exhaustive sampling of the entire state space.

In summary, the observed frequency of visits of different attractor states follows an approximate power law, but such behavior is most likely the consequence of sub-sampling of a distribution which is approximately log-normal.

## 3. Mean-field theory

We followed the methods of others (Ahmadian et al., 2015; Stern et al., 2014) to develop a mean-field theory for the large-*N* limit (*N* → ∞) of each system. The following description of the method has a slightly different emphasis from those of others, in part to connect to the alternative methods of section 4, and in part because our focus is on the existence of multiple stable fixed points rather than on the more general dynamics of the system.

In the large-*N* limit, because each individual connection strength scales to zero, the impact of small motifs (*e*.*g*., small subsets of units with net positive interactions) and correlations in activity between units becomes negligible. Therefore, the existence and stability of any state can be assessed by assuming all units receive input sampled from the same distribution arising from the sum of connection strengths multiplied by the activities of units. In small systems, the common scenario that units with positive connections are more likely to be coactive together, renders the simplifying assumption inaccurate. The large-*N* limit is also then (as stated in (Stern et al., 2014)) equivalent to averaging over all realizations of the connectivity matrix, *J*_*ij*_, which removes any correlation between individual units.

In the absence of any unit-specific identity, the unit label can be dropped from the formalism and the dynamical mean field equation is one for the distribution of activations represented by the variable ***x***(*t*) in the face of a distribution of inputs given by a new variable, ***η***(*t*), which we call the “field”:

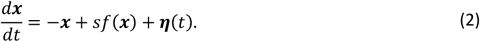

Self-consistency requires that the field, ***η***(*t*), is produced by the sum of the product of distribution of activities, *f*2***x***(*t*)3 (which result from the distribution of activation variable, ***x***(*t*)) multiplied by the connectivity matrix. Given the Central Limit Theorem, ***η*** is distributed as a Gaussian (a result justified more rigorously by others (Sompolinsky & Crisanti, 2018)) and given the lack of correlations between activity and connectivity in the large-*N* limit, we have:

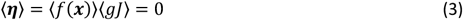

and

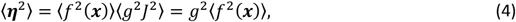

where we have used *J* to represent the *N* → ∞ limit of 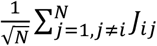.

Fixed points of the dynamics (Eq. 2) arise for the distribution of activations, ***x***, where

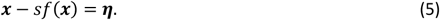

We define the variance of the zero-mean Gaussian distribution of ***η*** as σ^2^, such that

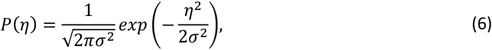

where σ^2^ must be calculated self-consistently from Eq. 4. For the system to possess multiple attractor states, the above set of equations (2)-(6) must have multiple solutions, and those solutions must correspond to stable states.

Multiple solutions arise from Eq. (5) if the function *x* − *sf*(*x*) is non-monotonic. Given the neural response function, *f*(*x*), has zero slope at very negative or very positive *x*, the function *x* − *sf*(*x*) has slope of +1 at these extremes and is therefore non-monotonic if for any value of *x* we have *f*′ (*x*) > 1/*s*. That is, multistability is possible if *s* > max *f*′(*x*). Hence the result in (Stern et al., 2014) that if *f*(*x*) = tanh(*x*) with a maximum gradient of 1, multistability is only possible if *s* > 1. For 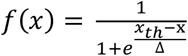 the requirement is *s* > 4Δ.

It is important to note that multiple self-consistent solutions of Eq. (4) for the variance of the field, ***η***, are also possible. Indeed, for the logistic input-output function, a solution with low variance corresponding to a quiescent, or low-activity state (in which activities of units are tightly clustered around *f*(0)) can coexist with a solution of greater input-variance. We assess both the stability of the solution with minimal activation (and therefore minimal variance of the field) as well as the existence of and stability of distinct solutions with higher activation when determining which states exist for a given set of parameters (see Appendix 3 for methods).

### 3.1. Phase diagram for networks with logistic single-unit response functions

In Figure 6, we show how the phase diagram depends on the slope of the input-output function, 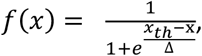, with the panels from A to F depicting results of increasing steepness (by lowering Δ). The final panel (Δ= 0) is produced by methods described in the next section. In all cases *x*_*th*_ is adjusted according to Eq. A1, to ensure single-unit bistability at *s* = 1. As can be seen, the minimum level of *s* allowing for multiple stable active states falls in proportion to Δ, and in the range 4Δ< *s* < 1, the system is quiescent at *g* = 0, but with increasing *g* becomes multistable. In all cases, and for all values of *s*, the expected transition to chaos arises with large enough *g*, though that transition is not always visible in the parameter ranges shown.

**Figure 6.**
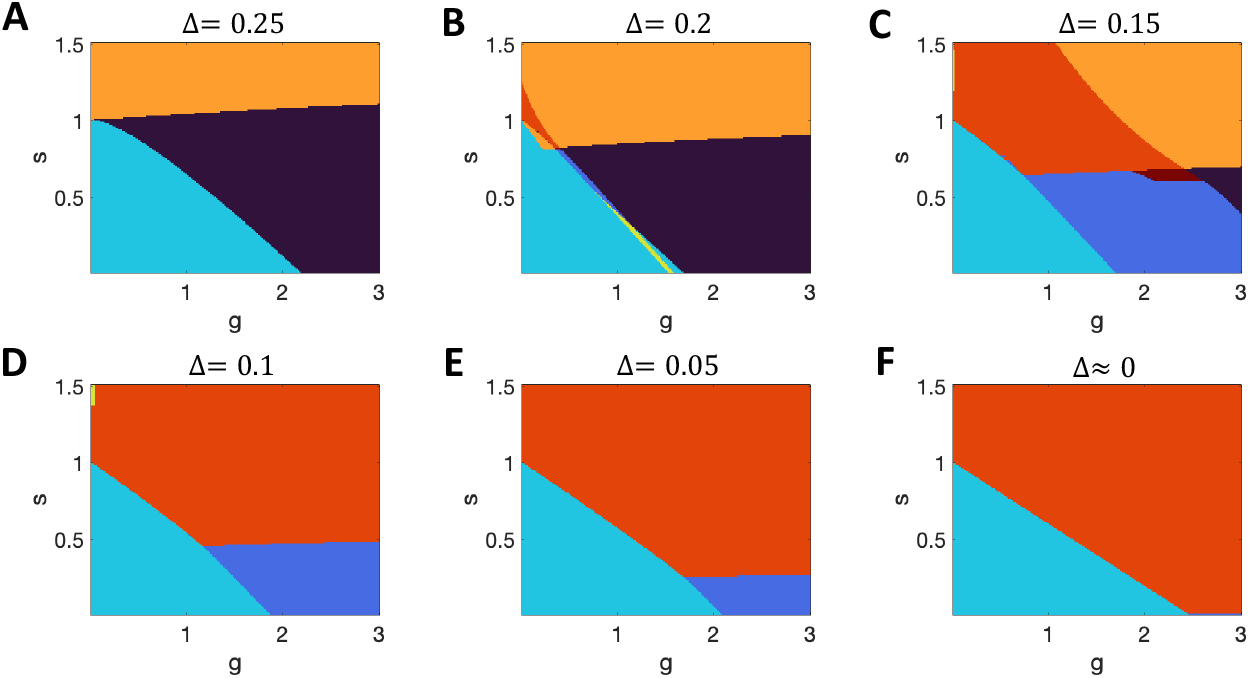
Multiple coexisting states in the infinite system with logistic input-output functions. **A-F**. Increasing steepness of the f-I curve is achieved by reducing Δ, with maximum slope 1/(4Δ). Key: black, chaos only; dark blue, quiescence + chaos; cyan, quiescence only; yellow, quiescence + single multi-unit active state; orange, multiple states all active; red, quiescence + multiple active states; crimson, chaos + multiple active states.

In systems with 0 < Δ< 0.25 (Figure 6 B-E) we find ranges of parameters for which the field, ***η***, does indeed have two self-consistent solutions. Most commonly, at low *s*, the cyan regions indicate the presence of a stable quiescent state with an unstable active state, that is the coexistence of inactivity and chaos in a given network. In a smaller range of parameters, the yellow region (Figure 6B) indicates the coexistence of a stable quiescent state with a stable active state. Such multistability exists even in the absence of cross-connections (*s* = 0) and concurs with our simulation results in Section 2.1. Indeed, the region of multistability spans the value of *g* = 1.55, observed at larger-*N* in simulations. Therefore, even without the self-excitation needed for individual units to be bistable with sufficient input, the network can possess multiple stable states, with the two distinct states resulting from and causing two distinct population-mean (and mean-square) firing rates and two distinct population input distributions.

3.1.1 Accounting for extreme tails of a Gaussian in the Infinite System

Our definition of the quiescent state contains a requirement that all units have activity of less than half of their maximum. In practice, in systems where bistability is possible (4Δ< *s* < 1) the quiescent state requires that all units are stable on the lower branch of the bifurcation curve. In the infinite system, any requirement of *all* units raises a subtle issue that we address in this subsection (and in Appendix C and Supplementary Figures 3 and 5).

The field, ***η***, which indicates the probability of any unit receiving a given input, follows a Gaussian distribution. When all firing rates are very low, the variance, σ^2^, of the Gaussian distribution for *η* is very low but is non-zero. The probability of a unit receiving input with a magnitude *Z*σ, that is many times, *Z*, greater than the standard deviation, σ, is vanishingly small (*e. g*. if *Z* = 6 the probability is less than 10^−9^ and if *Z* = 9 the probability is less than 10^−18^). However, for any finite *Z* the probability is strictly non-zero for any finite-level of input, so an infinite system will always have units whose inputs exceed that value. Therefore, in a system in which *s* > 4Δ, the quiescent state is unstable for an infinite system, unless *g* = 0 precisely. Yet, for any biologically feasible circuit we can define a *Z*_*max*_ and require the bifurcation points, *η*^∗^, to be within the range −*Z*_*max*_σ ≤ *η*^∗^ ≤ *Z*_*max*_σ, in order for the quiescent state to be defined as unstable. In this manner, we can study a system in which we have ignored correlations (an approach strictly only correct in the infinite-*N* limit) but at the same time define states that would be present in a large finite system with results accurate (to 1 part in 1000) for sizes up to *N* = 10^D^ (with *Z*_*max*_ = 6) or even *N* = 10^/E^ (with *Z*_*max*_ = 9).

A similar issue arises when we consider whether a system has multiple stable active states. The number of such states depends (exponentially) on the number of units receiving input between the two bifurcation points of ***x***(***η***) such that the unit could be either active or inactive for that level of input. Whenever there is a pair of bifurcation points (4Δ< *s* < 1), the argument from the previous paragraph again indicates that in an infinite system there is always a unit with input in that range. However, while in the infinite system the network is multistable, for any realistic system—even a large one—it may be very unlikely that any unit receives sufficiently extreme input, so we use the same value of *Z*_*max*_ to indicate multistability as we do for stability of the quiescent state.

Therefore, in Supplementary Figure 3, we replot the phase diagrams for distinct levels of *Z*_*max*_, while using a standard of *Z*_*max*_ = 6 for most phase diagrams. For example, with *Z*_*max*_ = 6, a large network of 10^D^ units would only have a probability of 0.001 of behaving differently from that indicated in the phase diagram (or a network of 10^3^ units would have a 1 in a million chance of behaving differently—the fewer the units in a network, the less likely at least one of those units receives excessively high input). In the limit of *Z*_*max*_ → ∞, the system has a discontinuity moving away from the y-axis, as even while the region of multistability approaches *g* = 0 (for 4Δ< *s* < 1) throughout this paper we have set the threshold, *x*_*th*_, such that disconnected units are only bistable if *s* > 1.

By contrast, when analyzing the network with tanh units of low Δ and higher *x*_*th*_—that is a steeper f-I curve, but with single-unit bifurcation maintained at *s* = 1—the output of units with zero input is close to -1, rather than 0. Such maximally negative output produces a larger variance of inputs across units, such that multistability is more common at very low cross-connection strength (Supplementary Figure 4). Therefore, while the choice of *Z*_*max*_ still impacts the phase diagram for those reasons discussed above, it does so to a much smaller extent for tanh units (Supplementary Figure 5).

### 3.2. Multistability without self-connections

Multistability without self-connections (*i*.*e*., with *s* = 0) is present in all networks with logistic response functions if we allow *x*_*th*_ to vary (or equivalently apply uniform input). To demonstrate this, in Figure 7A we show the phase diagram as a function of *x*_*th*_ and *g* for a system with *s* = 0 and Δ= 0.25—in this Case 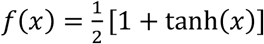. Systems with different Δ produce identical figures if the two axes are scaled by the same factor as Δ. As can be seen, the region of coexistence of quiescence with chaos is contiguous with and extends a region of coexistence of quiescence with an active stable state. These two regions, which depend on multiple stable solutions for the self-consistency of the field, ***η***, are not present if the response function is *f*(*x*) = tanh (*x*) (Figure 7B).

**Figure 7.**
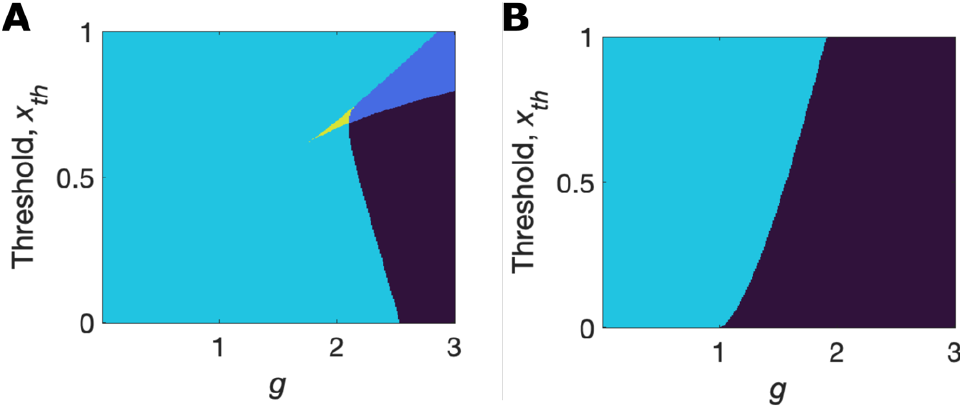
Phase diagram without self-connections. **A**. Network with logistic units does not require self-connections to be multistable. **B**. Network with tanh units is only monostable or chaotic without self-connections. (Black = chaotic; dark blue = chaotic + stable quiescence; cyan = stable quiescence only; yellow = stable quiescence + stable active state.)

The two response functions, 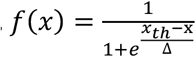 and 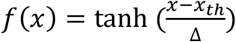, have a key difference that leads to them producing qualitatively different behavior. For the logistic function, the minimal absolute value of *f*(*x*) coincides with the minimal gradient of the function (if |*f*(*x*)| is low then *f*′(*x*) is low) whereas for the tanh function the opposite is true (if |*f*(*x*)| is low then *f*′(*x*) is near its maximum). Hence for the logistic function, it is possible for a narrow range of inputs to produce a stable narrow set of low firing rates (maintaining low inputs) while a solution with a large range of inputs leads to some much larger stable firing rates (maintaining high inputs) given the supralinearity of the response function. However, for the tanh function, the marginal feedback decreases with a change in rate from zero, so only one solution can exist.

## 4. Analysis of networks of units with binary response functions

For a system with binary units, *f*(*x*) = *Heaviside*(*x* − *x*_*th*_), the analysis simplifies, because a state is stable if all active units have input from other active units exceeding *x*_*th*_ − *s* and all inactive units have summed input from the active units less than *x*_*th*_. We define *k* as the number of active units, each with an activity of 1, so the above requirements on network inputs correspond to the sum of *k* − 1 of the connection strengths to each of *k* active units and to the sum of *k* connection strengths to each of the *N* − *k* inactive units.

Our main approximation is to treat these sums of connections strengths as independent draws from a Gaussian distribution with mean of zero and whose variance is 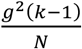 and 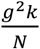 respectively. We then assume a solution with *k* active units exists if, given *N* independent draws from a Gaussian of unit variance, the top *k* draws, when multiplied by 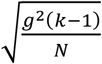 are greater than *x* − *s* (high input to active units) while the remaining *N* − *k* draws, when multiplied by 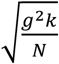 are less than *x*_*th*_. This is equivalent to the requirement that the (*k* + 1)_*th*_ greatest sample out of *N* samples, *X*_k+1,*N*_, from a unit-variance, zero-mean Gaussian distribution lies in the range:

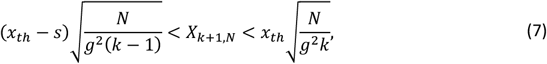

where we have assumed *x*_*th*_ − *s* > 0 and *x*_*th*_ > 0 (which holds in our standard system with *x*_*th*_ = 1 so long as *s* < 1, and which is the parameter region in which we have greatest interest).

Such a requirement can be calculated using the methods of order statistics (David & Nagaraja, 2003), which we follow for the Gaussian distribution, defining

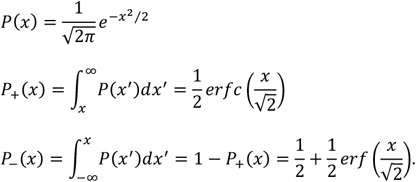

such that

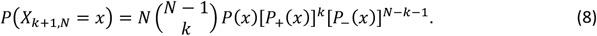

Therefore, we have a stable state with *k* of *N* units active with a probability *P*(*k, N*) given by:

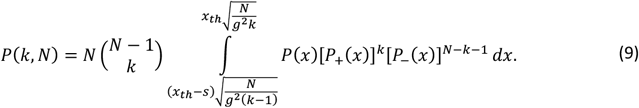

We then calculate the probability a system has at least one stable state with multiple active units, and so is multistable (as the quiescent state is always stable for *x*_*th*_ > 0) for a given system size, *N*, via:

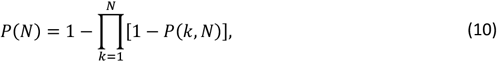

(there is only an absence of multistability if it is absent for all possible *k*). Again, (10) is an approximation as it assumes *P*(*k, N*) is independent for different values of *k*. We will see that in the large-*N* limit the approximation becomes exact, as *P*(*k, N*) becomes either 0 or 1, so that *P*(*N*) is also either 0 or 1, and we have multistability with probability 1 if and only if it arises with probability 1 for some value of *k*, that is *P*(*N*) = max *P*(*k, N*).

### 4.1. Finite-*N* results of analysis with binary units

Our simulation results presented in Section 2 (Figure 4) suggest that, for some parameters, networks of intermediate size have the greatest probability of multistability. Given the increasing likelihood of missing stable states as *N* increases using simulation methods, that simulation result may be incorrect due to undersampling at large *N*. Therefore, we use our approximate analytical methods for networks with binary units to address the *N*-dependence of the probability of multistability, by solving Equations (8)-(10) above.

In Figure 8A-C, we show that for small networks of *N* = 6, *N* = 12, and *N* = 18, for which we can exhaustively test all initial conditions and therefore find all stable states in simulations, the approximate analytical method (solid line, which plots Eq. 9) compares well with the simulated data (crosses). Moreover, in Figure 8D-F, when we use Eq. 10 (red lines) to estimate the probability the network is multistable, the simulated results (crosses) are remarkably close to the analytic approximation. Such a result is surprising, as one would expect a positive correlation across networks and the numbers of stable states. The blue lines in Figure 8D-F are the results for a correlation of +1, in which the network’s probability of multistability is simply the maximum across possible states and is much farther from the data than the analysis assuming zero correlation (the red line). Nevertheless, across all methods, in Figure 8D-E, we do indeed find that the probability of multistability peaks at intermediate network size, remarkably reaching values of approximately 1 for *s* = 0.5, *g* = 1.2, before falling to zero at large network size (*N* > 30000).

**Figure 8.**
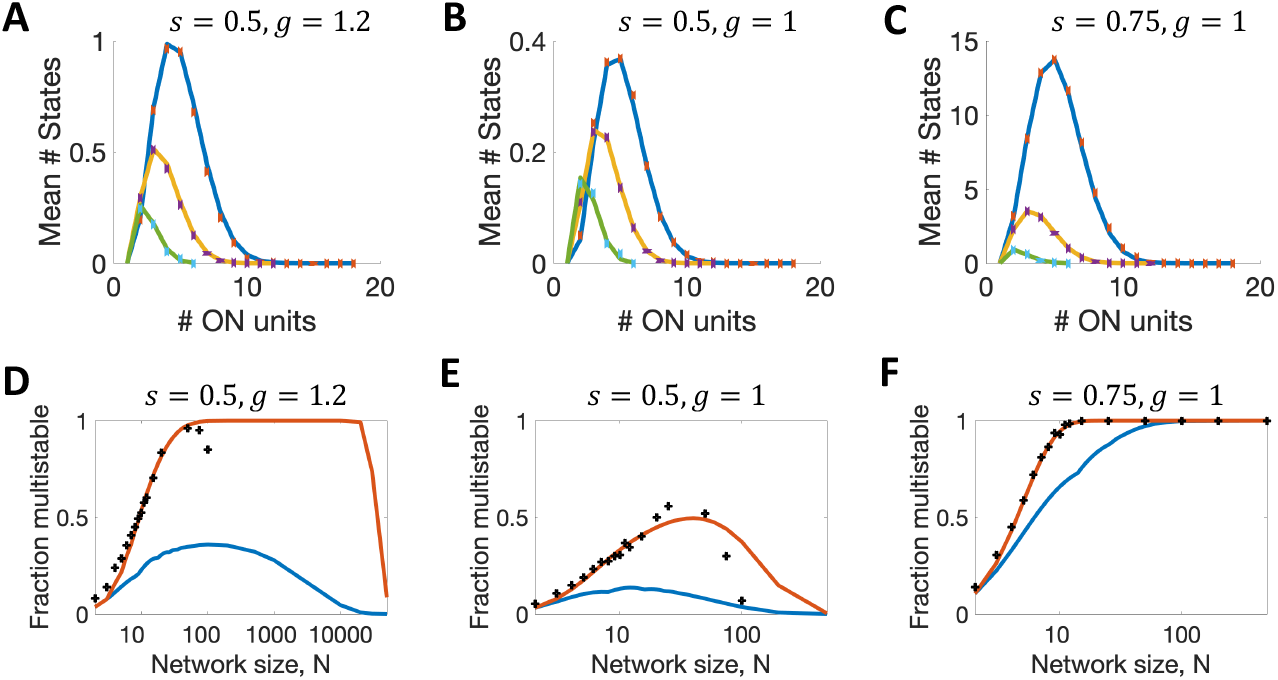
Numbers of stable states in finite binary networks. **A-C**. Expected number of stable states with *k* active units when *N* = 6 (green), *N* = 12 (yellow), *N* = 18, blue. Continuous lines from Eq. 9, points from simulations. **D-F**. Probability a network is multistable as a function of network size, *N*. Top curve uses Eq. 10, lower curve uses max *P*(*k, N*). Crosses are simulated data points in all panels, and are an undercount at large *N*.

### 4.2. Large-*N* limit of system with binary units

In the large-*N* limit, the above equations (8)-(9) can be simplified. First, we note that

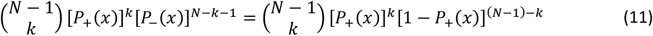

is the Binomial probability for achieving *k* outcomes from *N* − 1 independent selections, with individual probability of outcome, *P*_+_(*x*). In general, the probability has a peak at the integer value of *k* closest to (*N* − 1)*P*_+_(*x*), with a standard deviation of 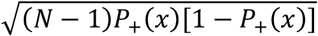.

If we define 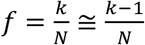 with *k* ≫ 1 as well as *N* ≫ 1, then the integration limits for *P*(*k, N*) become 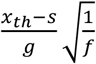 and 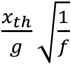. Also, for large *N*, the Binomial probability term approaches a Dirac delta-function at the value of *f* = *P*_+_(*x*), as it’s standard deviation in *f* scales as 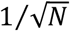.

Therefore, in the large-*N* limit, *P*(*f*) = 1 if the integration range over *x* contains the value where *f* = *P*_+_(*x*) and *P*(*f*) = 0 otherwise. Algebraically this becomes a requirement that *f* lies between two thresholds, Θ_1_ and Θ_2_, which each depend on *f*:

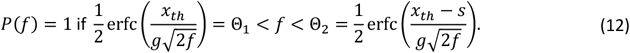

Figure 9A indicates the region of these inequalities as a function of *f* for the specific values of *x*_*th*_ = 1, *s* = 0.5, and *g* = 2.75, with Figure 9B showing that for a wide range of *g* the possible numbers of active units in a stable state splits into two distinct ranges. Simulation results in Figure 9C demonstrate such bimodality in numbers of active units for a similar network (*s* = 0.75 and *g* = 2.5) with *N* = 400.

**Figure 9.**
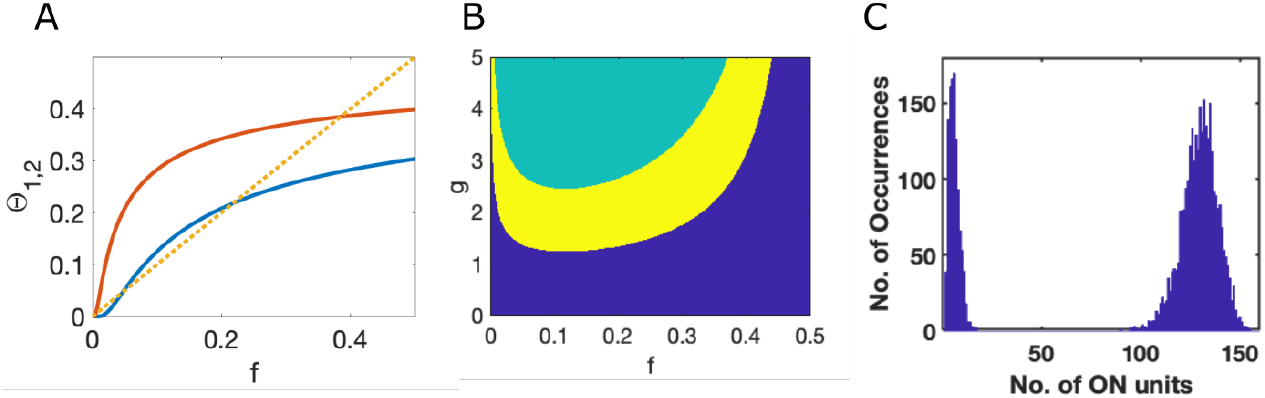
Number of active units in stable states is bimodal. **A**. The two complementary error functions producing the bounds on fraction of active units, *f*, from Eq. (12) (infinite system with Δ= 0, *x*_*th*_ = 1, *s* = 0.5, and *g* = 2.75). Where the red curve is above the dashed yellow line active units are stably active, but otherwise not. Where the blue curve is below the dashed yellow line, inactive units are stably inactive, but otherwise not. Note the two distinct ranges of *f* in whch the dashed yellow line is above the blue line and below the red line. **B**. Across a range of *g* for the same system as **A**, the allowed range of *f* for which stable solutions are possible, as indicated in yellow, splits in two. In the blue region the active units are unstable, while in the green region the inactive units are unstable. **C**. Simulations of a network (Δ= 0, *x*_*th*_ = 1, *s* = 0.75, and *g* = 2.5) of size *N* = 400 from 10^5^ initial conditions of varying numbers of active units lead to final stable states with a bimodal distribution in the number of active units.

Given that at 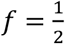 it is always true that 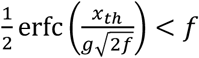 (because of the limited range of the complementary error function) the criterion for Eq. 12 to have a solution for some value of *f* is the requirement 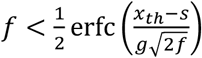 for some *f*, which leads to a minimum value of *g* = *g*^∗^ at which the lines *y* = *x* and 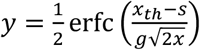 meet at a tangent. The critical value occurs where 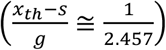 such that, as a function of *s* and with *x*_*th*_ = 1, multistability arises in this system if *g* > *g*^∗^(*s*) ≅ 2.457(1 − *s*) (Figure 6F).

Notice that as *s* approaches zero, the range of *f* with allowed solutions shrinks toward the line Where 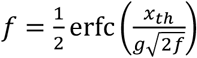. For *g* > *g*^∗^, there are two such solutions, which are distinct crossings of the line. The distinct solutions indicate two separate ranges for the possible number of active units in stable attractors. In the example shown in Figure 9B, solutions are possible in two ranges, either with a very low fraction (<5%) of units active or with a fraction in the range of 30%-40% of units active. The two distinct ranges are also visible from simulations as a bimodal distribution in the numbers of active units in stable states following random initial conditions (Figure 9C).

A lower bound on the fraction of active units arises, because with few active units there is too little network input to activate those units. The upper bound arises because half of the units receive net negative input, so cannot be stably active if *s* < 1, and only a subset of those units receiving net positive input receive an amount greater than 1 − *s*, as needed to be stably active. The bounded region in which active units have sufficient input to remain stably active can contain within it a separate bounded region of instability (Figure 9) because the random network input can be sufficiently strong that some of the inactive units (of which there are more than there are active units) receive too much input to remain “off”.

## 5. Discussion

Firing rate models of neurons are valuable because they represent the likely states of a neural circuit in a relatively simple manner and can be solved rapidly. The foundation of a firing rate model is the input-output function of a neuron, which is typically designed to have bounded outputs over the domain of inputs. For its ease of mathematical manipulation, the hyperbolic tangent function, *f*(*x*) = tanh(*x*), has been used with great success, most notably for first demonstrating the transition from quiescence to chaos as the strength of random cross-connections increases (Sompolinsky et al., 1988; Stern et al., 2014). The negative portion of tanh(*x*), while it cannot correspond to negative firing rates, could be considered representative of a group of mixed excitatory-inhibitory neurons in which the mean rate of inhibitory neurons exceeds that of excitatory neurons.

Given the function *f*(*x*) = tanh(*x*) is simply a translated version of the function *f*(*x*) = tanh(*x* − *x*_*th*_) + 1, one might expect that analysis of a system with units responding via the one function would provide all the qualitive insight necessary to understand the behavior of a system with units responding via the other function. However, this is not the case. A disconnect between the behavior of a system of neurons with *f*(*x*) = tanh(*x*) and that of a system with *f*(*x*) = tanh(*x*) + 1 has been shown by others (Figure 4b of (Touboul & Ermentrout, 2011)) whereby a Hopf bifurcation disappears as the input-output function of neurons is parametrically shifted up toward non-negative values. In our analyses, we find two qualitative changes. The first is a shift in phase boundaries leading to the result that random cross-connections, whose mean value is zero, can produce multistability in a system in which single units are not in of themselves bistable.

Second, we find the possibility of bistability via distinct stable solutions for the self-consistency of the field. Alternative self-consistent solutions of the field can lead to multistability arising from random, zero-mean, cross-connections even in systems without self-connections (Figure 3 and Figure 6B). The distinct self-consistent field solutions, with different variances in the input currents, correspond to states with distinctly different numbers of active units. Figure 9 indicates a similar bimodality in the numbers of active units in simulated binary-unit systems and is coupled with an analysis of how such bimodality arises in the system.

We find a subtlety when taking the infinite limit of our system using the logistic input-output function, with a strict discontinuity between results with *g* = 0 and those with *g* = *ϵ* (where *g* scales the strength of cross-connections and *ϵ* is an infinitesimal positive quantity). The reason being that for non-zero *g*, there is a non-zero (even if miniscule) probability that the within-circuit input to a unit, which is drawn from a Gaussian with width proportional to *g*, is sufficiently strong to render that unit bistable. However, when the bifurcation point is many tens of standard deviations above the zero mean of the Gaussian distribution, the probability becomes infinitesimal and is irrelevant in any real or simulated system, even with billions of units. For similar reasons, the strict mathematical limit has a discontinuity when altering the width of the logistic function from Δ= 0 to Δ= *ϵ*. If, instead of producing a phase diagram, with a sharp boundary for multistability, we focused on the entropy of the system (the log of the number of stable states) scaled by system size, *N*, such discontinuities would disappear as the entropy would reduce continuously and smoothly (and rapidly) from the boundaries of multistability shown in Figure 6, to a tiny value before becoming strictly zero at *g* = 0 or Δ= 0.

Multistability, when exhibited as a set of discrete stable fixed points, may seem unlikely in any cortical circuit given that activity is never static *in vivo*. However, a network based on multiple fixed points, but with randomly timed transitions between them, can match the observed data in a number of systems (Ballintyn et al., 2019; Ksander et al., 2021; La Camera et al., 2019; Mazzucato et al., 2019; Miller, 2016; Miller & Katz, 2010; Moreno-Bote et al., 2007; Recanatesis et al., 2022). Moreover, analyses of patterns of neural spiking *in vivo* have, in many cases, shown that a discrete state-based formalism better matches the data than a formalism assuming continuously changing, graded activity (Abeles et al., 1995; Miller & Katz, 2010, 2011; Ponce-Alvarez et al., 2012; Sadacca et al., 2016; Seidemann et al., 1996).

While the strengths of connections between units are treated as independent random variables for ease of analysis in this paper, in practice there is internal structure in the connectivity among neurons, even between excitatory pyramidal cells (Song et al., 2005; Stepanyants & Chklovskii, 2005).

Moreover, connections from cortical neurons typically have fixed sign (all excitatory or all inhibitory) according to neuron class, a feature that can change the behavior of random networks (Rajan & Abbott, 2006). In our work, we consider a firing rate model unit as representing the mean rate of a cluster of many neurons (as is necessary to omit the pulsatile spike interaction from simulations) so the net interaction between units can be of either sign according to whether the dominant connections are excitatory-to-excitatory, or excitatory-to-inhibitory, etc. Moreover, much of the nonrandom cortical structure can be accounted for by considering the intra-cluster connectivity to be distinct from the inter-cluster connectivity (Bourjaily & Miller, 2011) as we do here.

Our main conclusion is that multistability can be produced via random, zero-mean cross-connections in neural circuits without the exceptionally strong self-connections needed to produce bistability in a single cluster of neurons (a unit in a firing-rate model) so long as the neurons without input have a low firing rate and if rate increases supralinearly with low input.

## Code Availability

MATLAB codes used to produce the results in this paper are available for public download at https://github.com/primon23/Multistability-Paper.

## Acknowledgments

The authors are grateful to NIH-NINDS for support of this work via R01 NS104818 and to the Swartz Foundation for a fellowship to SQ. SM is grateful to Merav Stern for helpful conversations in the early stages of this work.

## Statements and Declarations

The work was supported by a grant from the National Institutes of Health, R01 NS104818, by the Swartz Foundation, and by the Neuroscience Graduate Program of Brandeis University.

## Author Contributions

All authors contributed to the writing and editing of the manuscript and approved the final version. Simulations were carried out by Jordan Breffle, analysis by Subhadra Mokhashe, Siwei Qiu, and Paul Miller. The first draft was written by Paul Miller, who also conceived of the project.

## Appendix 1: Monte Carlo simulation method

Our standard procedure was to simulate 100 different realizations of the connectivity matrix to produce 100 random networks for a given parameter combination. For each connectivity matrix, we then completed sets of multiple trials, each trial with a distinct initial condition (100 trials for perturbation analysis in Figure 2 and for scaling *g* in Figure 3; 200 trials for parameter grids in Figure 4; and 10^6^ or 10^5^ trials respectively for the networks with binary units in Figures 8 and 9). For the small (*N* ≤ 25) networks with binary units in Figure 8 all 2^*N*^combinations of initial conditions were used with each unit at an initial rate of its minimum or maximum.

The continuous models were simulated using MATLAB’s ode45 function. Each trial was simulated until either a maximum simulation time was reached (5,000 *τ* for Figure 3 and 10,000 *τ* for Figure 4), or until a stopping condition was reached in the case that the maximum 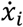 at a give timestep was less than 2 ∗ 10^−D^. If this stopping condition was reached, then the activity was considered to have reached a stable state because the network possessed a point attractor at that set of firing rates.

Logistic units were classified as active if their firing rate exceeded 0.5. Tanh units were considered active if the absolute value of their rate exceeded 0.001. For the continuous models, typically the first trial was initialized with inputs near zero, to test if the quiescent state was stable. For all subsequent trials, the initial rates of the units were set to a uniform random distribution over 0 to 1 and transformed by a logistic function with *x*_*th*_ = 0.5 and Δ= 0.1.

For the perturbation analysis, each trial of each network was simulated for a full 21,000 *τ*. Then, at each of 100 linearly spaced time points between 20,000 and 20,800 *τ* 10% of the units’ firing rates were randomly perturbed upwards or downwards by 10^−5^ and the simulation was then continued from each such perturbed state for 200 *τ*. The root mean squared (RMS) deviation of the perturbed simulation from the original simulation quantified the extent to which the perturbation caused a divergence in activity. The median RMS deviation over the 100 perturbations was then used to classify each trial as a point attractor, a limit cycle, or chaotic. The median RMS deviation exponentially decayed for point attractors, exponentially increased for chaos, and increased but reached a plateau at a low level for limit cycles. Classification thresholds were set based on the R^2^ of a linear fit to the exponential RMS deviation and the magnitude of the RMS deviation averaged between 190 to 200 *τ* post-perturbation. Trials with final RMS deviations below half the magnitude of the initial perturbation and with no units having a change in their firing rate exceeding 10^−4^ in the last 10 *τ* of the unperturbed simulation were classified as point attractors. To classify trials as chaotic vs limit cycles, a classification boundary was determined as a function of each trials’ linear fit R^2^ and final RMS deviation. Trials above the line 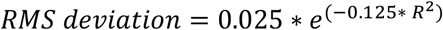 were classified as chaotic. This boundary allows the separation between these two dynamics because it accounted for both chaotic trials that very quickly converged to a large RMS deviation (large RMS deviation and low R^2^) and chaotic trials that had a slower exponential increase in their RMS deviation (lower RMS deviation at 190 to 200 *τ* but high R^2^). Final activity states of the unperturbed simulations were used to confirm these classifications.

## Appendix 2: Choice of single-unit input threshold

For comparison across systems with distinct single-unit input-output functions, *f*(*x*), we adjust the offset, *x*_*th*_, such that a single unit becomes bistable with self-connection strength of *s* = 1, in all cases.

For the logistic function, such a requirement means that a saddle-node bifurcation occurs at *s* = 1, with unstable and stable fixed points colliding at *x*^∗^ given by −*x*^∗^ + *sf*(*x*^∗^) = 0 such that *x*^∗^ = *f*(*x*^∗^) and 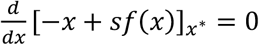 such that 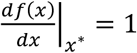. Combining these equations and using the result for the logistic function that 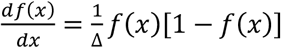 leads to the requirement:

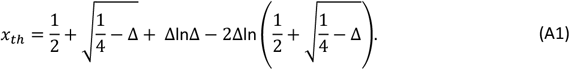

For the binary response function, *f*(*x*) = *Heaviside*(*x* − *x*_*th*_), we have *x*_*th*_ = 1, which can be seen from the above equation in the limit Δ→ 0.

For the hyperbolic tangent function, 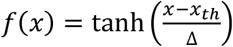, a similar derivation leads to

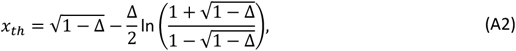

which yields *x*_*th*_ = 0 if Δ= 1, matching the simplest anti-symmetric response function, *f*(*x*) = tanh(*x*), and as with the binary response function, *x*_*th*_ = 1 if Δ= 0.

## Appendix 3: General mean-field methods

To test whether a distribution of the interacting variables, ***x***, produces a stable fixed point, it is necessary to obtain information about the eigenvalues of the Jacobian matrix of the dynamical equations expanded linearly about the fixed point (Strogatz, 2015). If all such eigenvalues have a negative real part then the fixed point is stable. Linearization around a fixed point, ***x***^∗^, yields

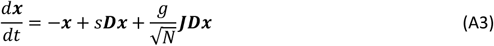

where ***D*** is a diagonal matrix with elements equal to the corresponding derivatives of the input-output function, *f*′(***x***^∗^), and ***J*** is the unit variance, zero mean, Gaussian connectivity matrix.

We follow the methods of others (Ahmadian et al., 2015; Stern et al., 2014) who showed that eigenvalues of such a system are found at the complex values, *z*, where

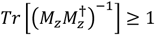

With

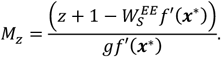

In the large-*N* limit the sum within the Trace become an integral over the distribution of activations, ***x***, to yield the criterion (Ahmadian et al., 2015; Stern et al., 2014):

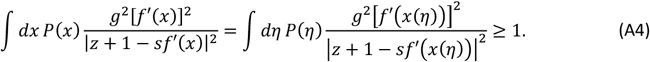

As noted by (Stern et al., 2014), for the system to be stable we require that Equation A4 is not satisfied for any *z* with *Re*[*z*] > 0, which allows us to assess the case where *Re*[*z*] = 0 and note that any non-zero contribution to *Im*[*z*] increases the absolute value of the denominator above, so if there are no eigenvalues with *z* = 0 there cannot be any on the imaginary axis. Therefore, in general we require, for there to be no eigenvalues with positive real part that

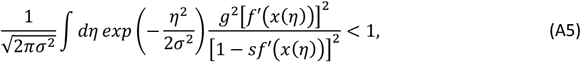

where we have substituted for *P*(*η*) and σ^2^ = *g*^2^⟨*f*^2^(*x*)⟩. We have also assumed that the function in the denominator, 1 − *sf*′2*x*(*η*)3, is positive, as any negative portion of the function means there is a divergent positive contribution to the integral for some *z* with *Re*[*z*] = *sf*′2*x*(*η*)3 − 1 > 0.

We are interested in cases of multistability, where the activations, *x*(*η*), can have more than one value based on the solutions *x* − *sf*(*x*) = *η* for some values of *η*. This requires that *x* − *sf*(*x*) is a non-monotonic function, which occurs if max [*f*′(*x*)] > 1/*s* (to produce a region of negative slope in the function *x* − *sf*(*x*)). The need for a region of negative slope arises because in all cases considered here at large positive or negative values of *x, f*^′^(*x*) = 0 and *x* − *sf*(*x*) has a slope of +1. In cases of multiple solutions for *x*(*η*), care must be taken in the choice of *x*(*η*), as while stability is enhanced by choosing the solution with the lower value of *f*′2*x*(*η*)3, such a choice can lead to the lower value of *f*^2^(*x*) for some input-output functions (but not if *f*(*x*) = tanh (*x*)) which can lead to the self-consistent solution for the distribution of *η* to become too narrow to support multistability, as discussed below.

In the logistic networks, 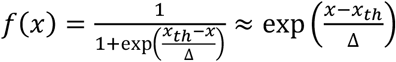 for *x* ≪ *x*, is never exactly zero. Therefore the Gaussian distribution of *η* will always have non-zero variance for *g* > 0 and, even if the distribution is narrow with very small variance, the distribution always retains some vanishingly small but non-zero density at the values of *η* required to support multiple solutions of *x*(*η*) if *s* > 4Δ. However, if bifurcation points in *x*(*η*) require levels of the Gaussian-distributed *η* that are many standard deviations from its mean of zero, such solutions give exponentially small probability of multistability in a finite network, so are unlikely to be observed in practice. Therefore, we set a threshold, *Z*_*max*_σ, in terms of the number, *Z*_*max*_, of standard deviations, σ, of the field of inputs, *η*, such that if both bifurcation points, *η*^∗^, are beyond the threshold (*η*^∗^ < −*Z*_*max*_ or *η*^∗^ > *Z*_*max*_) we ignore both the extra solutions and any instability they cause. To clarify the result of such a limit, we show results with multiple values of *Z*_*max*_ in Supplemental Figure 3 and Supplemental Figure 5, while using a default value of *Z*_*max*_ = 6 in other figures. In this manner, we have used the results for an infinite system in which correlations are absent, but applied them to a system in which the number of units could range from 10^3^ to 10^6^ to 10^15^ (as *Z*_*max*_ changes from 3 to 6 to 9) and the results be accurate for 999 networks in 1000 of that size. For further explanation see also the text in Section 3.1.

## Supplemental figures

**Supplementary Figure 1:**
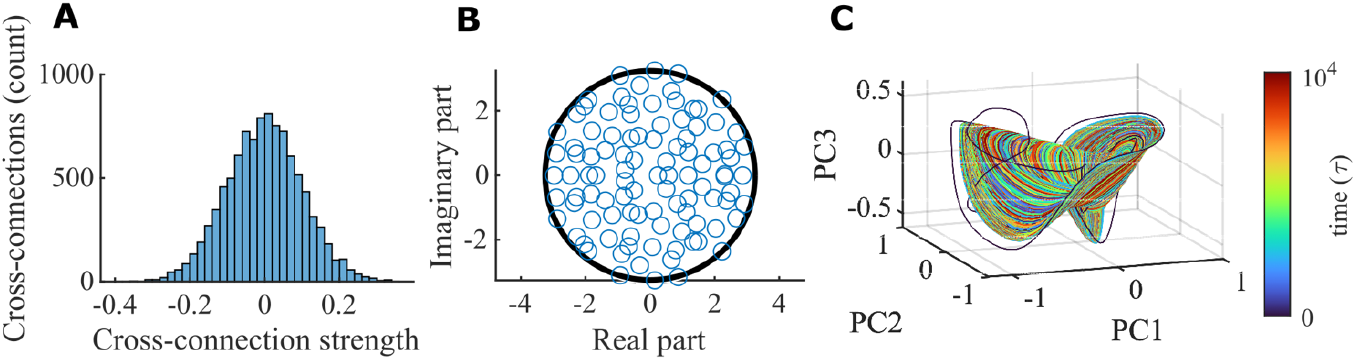
The example network in Fig. 2 is a typical random network. **A**. Distribution of cross connections of the example network in Figure 2. The cross-connection distribution has a mean of -0.00053 and standard deviation of 0.10037, which is consistent with the expected mean of 0 and standard deviation of 0.1 (KS-test, p = 0.708, KS-statistic = 0.007). **B**. Eigenvalues of the cross-connection matrix. The black circle shows the expected bounds for the large-N limit. **C**. The first three PCs of the chaotic attractor shown in yellow in Figure 2B. Color is simulation time. There is an initial transient that quickly converges to the chaotic attractor. Analysis of simulated perturbations shows that it is chaotic (Figure 2D).

**Supplementary Figure 2:**
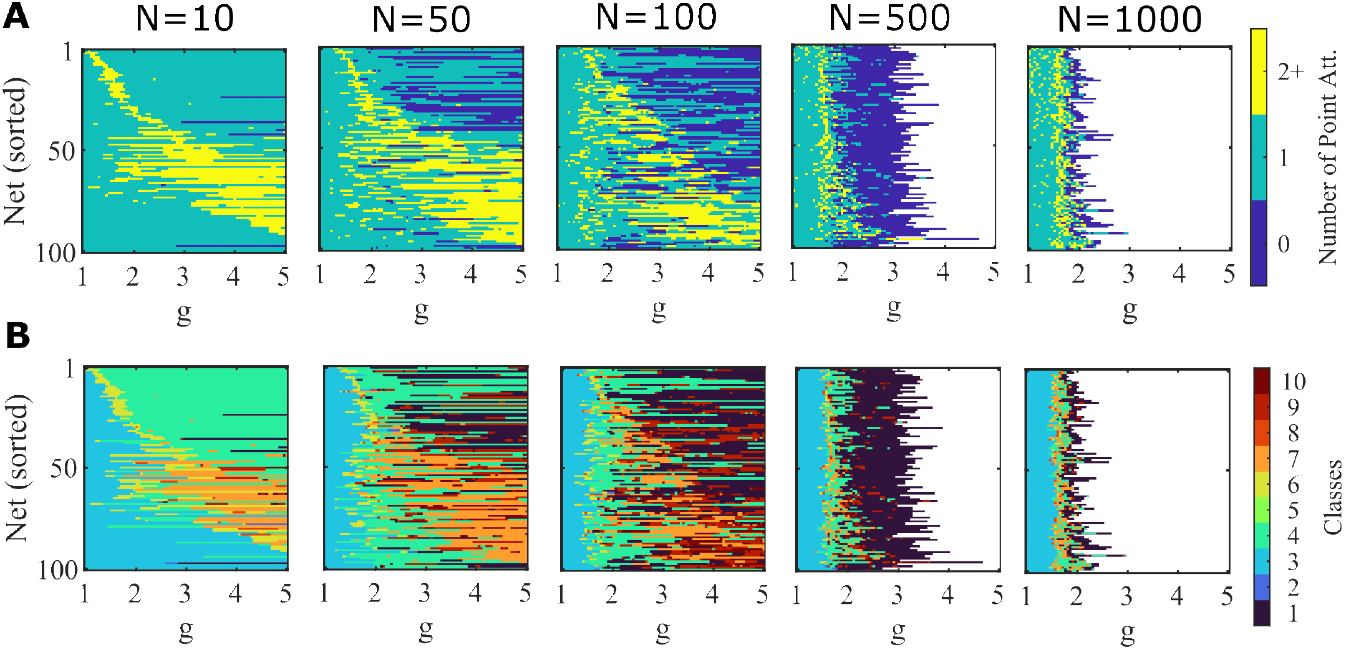
The dynamic regimes of individual networks changes as g is scaled. **A**. Similar to Figure 3B, but for networks simulated over a smaller range of g-values. 100 random networks of logistic units (Δ=0.2) with no self-connections (s=0) of varying size (N=10, 50, 100, 500, 1000) were simulated across g values. The same network can gain and lose mulistability as g is scaled. Color scale indicates number of point attractors found within 100 trials. White indicates g values that were not simulated due to computational limits. **B**. The same simulations as in **A**, but where color now indicates the classification of the set of activity observed across the 100 simulated trials. 1, no trials converge; 2, stable quiescence + some trials fail to converge; 3, only stable quiescence; 4, only a single stable active state; 5, some trials don’t converge + stable quiescence + at least one stable active state; 6, stable quiescence + a single stable active state; 7, multiple stable active states + no stable quiescence; 8, multiple stable active states + stable quiescence; 9, some trials fail to converge + a single stable active states; 10, some trials don’t converge + multistable.

**Supplementary Figure 3.**
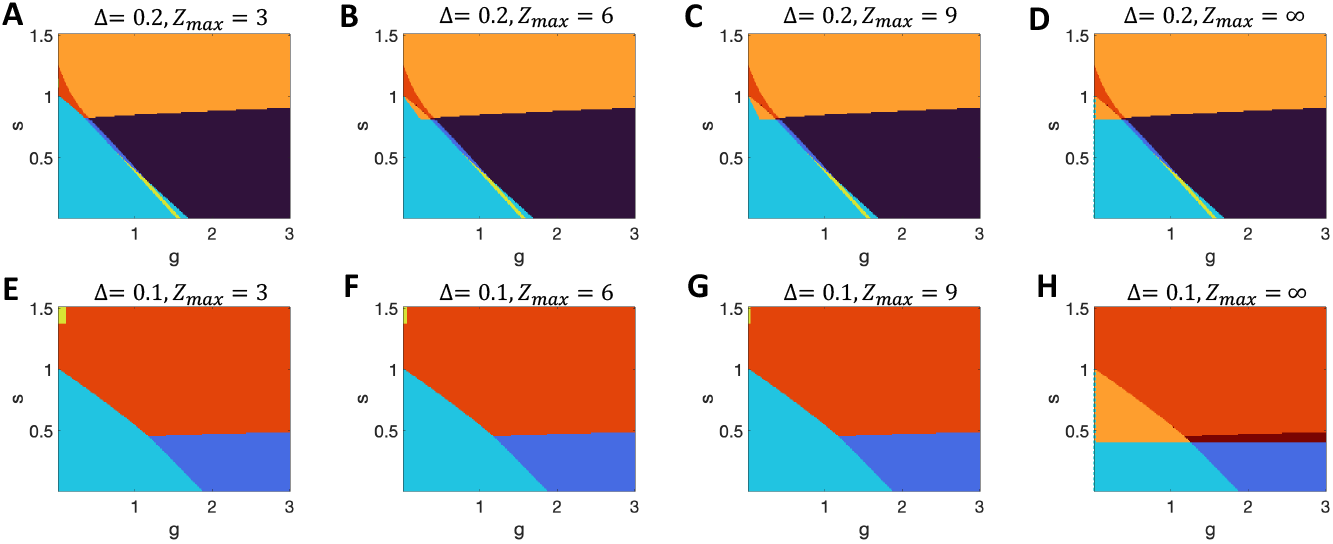
Impact of the criterion for multistability in networks with logistic units. **A-D** Results with Δ= 0.2 with varying threshold, *Z*_*max*_, for the number of standard deviations from the mean input that a unit must receive before considering a unit in the network to switch state. The mathematical limit is shown in **D**, while **A-C** indicate the multistable region growing with increased *Z*_*max*_. Note that in all cases the system with *g* = 0 on the y-axis can not be multistable for 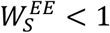. **E-H** Results for Δ= 0.1. (Black = chaos; dark blue = chaos + quiescent stable; cyan = quiescent only; yellow = quiescent + active stable state; orange = multiple active stable states; red = stable quiescent + multiple active stables states; crimson = chaos + multiple stable active states.)

**Supplementary Figure 4.**
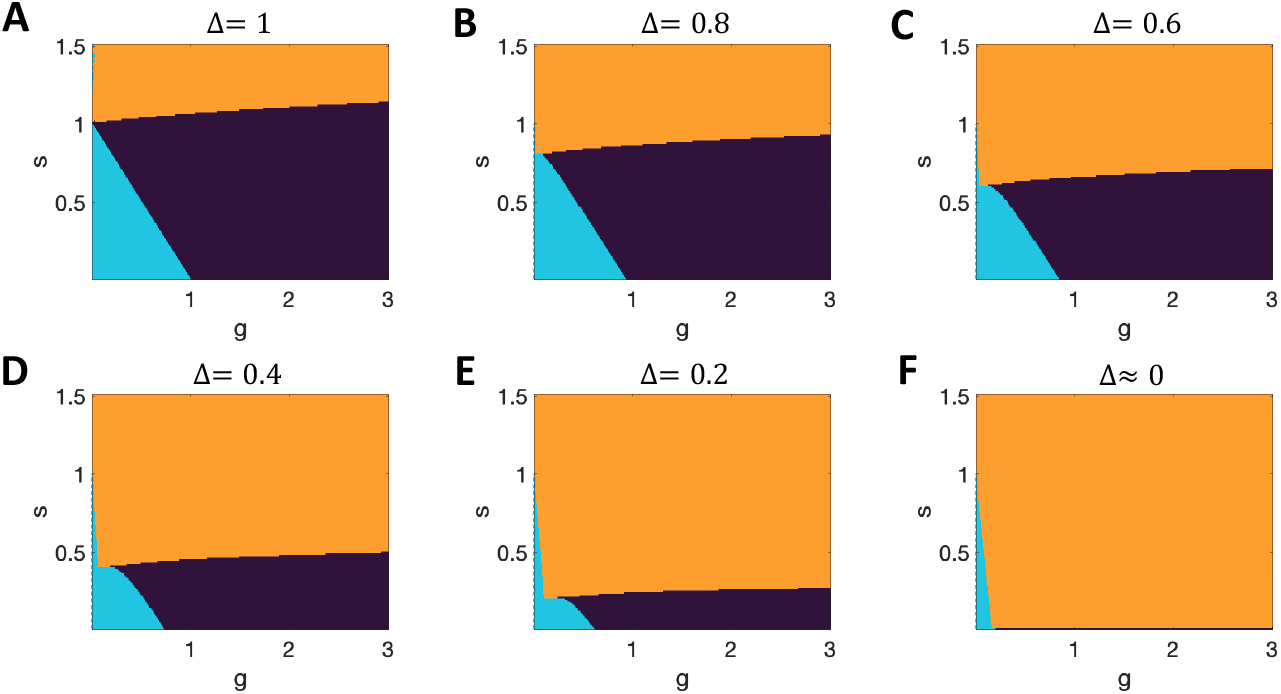
Phase diagram for networks with tanh units. **A**. Results with Δ= 1 replicate those of (Stern et al., 2014). **B-D**. Region of multistability increases to lower *s* while remaining only for *s* ≥ 1 on the y-axis. (Black = chaos; cyan = quiescent only; orange = multiple active stable states.)

**Supplementary Figure 5.**
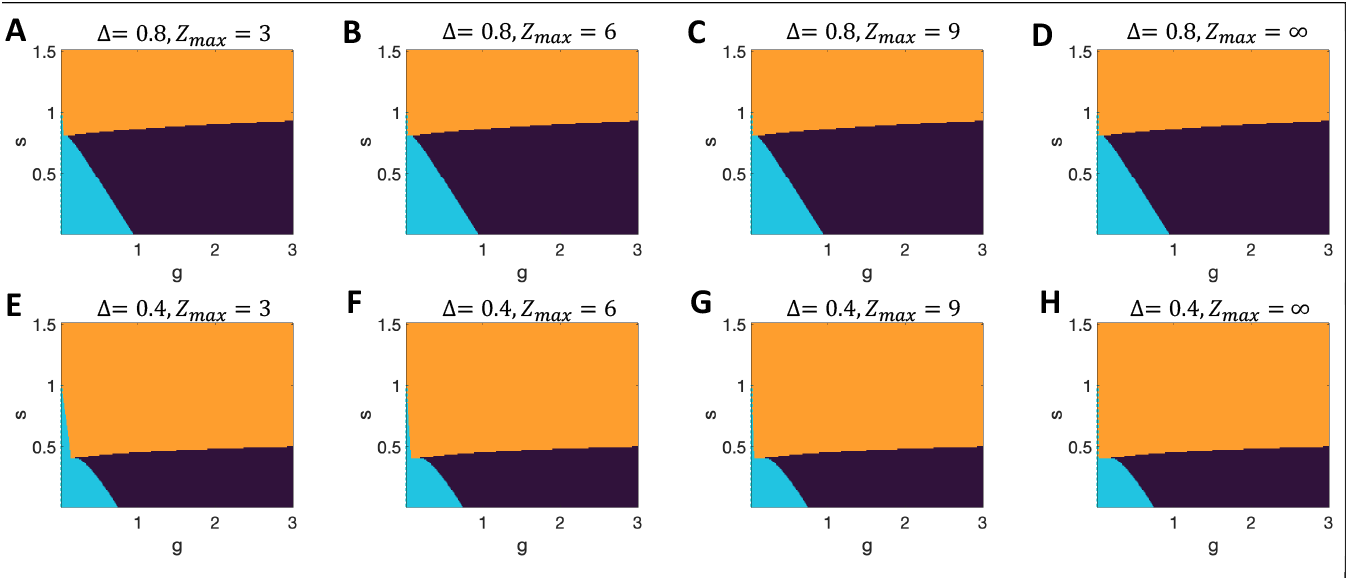
Impact of the criterion for multistability in networks with tanh units. **A-D**. Results with Δ= 0.8 with varying threshold, *Z*_*max*_, for the number of standard deviations from the mean input that a unit must receive before considering a unit in the network to switch state. The mathematical limit is shown in **D**, which in this case is minimally different from the results with lower *Z*_*max*_ in (**A-C)**. Note that in all cases the system with *g* = 0 on the y-axis can not be multistable for *s* < 1. **E-H**. Equivalent results for Δ= 0.4, with a tiny, but observable dependence on *Z*_*max*_. (Black = chaos; cyan = quiescent only; orange = multiple active stable states.)

